# One Health Approach for the sampling of different bat species living in a sympatric colony

**DOI:** 10.1101/2022.09.22.508887

**Authors:** Thejanee Perera, Sahan Siriwardana, Therese Muzeniek, Beate Becker-Ziaja, Dilara Bas, Fatimanur Bayram, Mizgin Öruc, Inoka Perera, Jagathpriya Weerasena, Shiroma Handunnetti, Franziska Schwarz, Gayani Premawansa, Sunil Premawansa, Wipula Yapa, Andreas Nitsche, Claudia Kohl

## Abstract

Bats are important contributors to the global ecosystems; at the same time, they are known to be a natural reservoir host for a number of human pathogenic viruses. These and many other unique features make them an interdisciplinary research object in the context of One Health, comprising zoology, ecology, virology, microbiology, molecular biology, immunology and public health issues. Performing field studies for bat research often aims to cover several of these topics and requires the combination of specific expertise in different fields. We carried out three individual field studies in Wavul Galge cave (Koslanda, Sri Lanka), where several bat species roost sympatrically. The main goals were to study the bat colony for ecological aspects and to sample bats for virological and molecular biological analyses. In the course of the field studies, we optimized the sampling procedure regarding safety aspects, a preferably low impact on the captured bats and an improved output of high-quality samples for further analysis. Different sampling methods and procedures were compared in order to establish a suitable strategy for frequent sampling and monitoring of these bats. In the present case study, we report on this process of optimizing our field work and provide suggestions for bat sampling methods that cause comparably less stress for the captured animals. We also report on constraints and obstacles encountered during the practical implementation and possible measures to overcome these.

With these practical experiences, we hope to give support to other interdisciplinary research teams preparing for bat field work. Furthermore, we emphasize the need for the respectful treatment of the animals and minimized disturbance of their natural habitat when carrying out sustainable bat research.

## Introduction

Bats of the order *Chiroptera* are mammals with the second-highest species diversity after rodents and are globally distributed [1]. Unique features like their ability to fly, migratory habits and longevity make them a promising topic in zoological and ecological research [2]. In addition, bats are known to be the natural reservoir hosts of numerous pathogens including zoonotic viruses causing diseases in humans [3].

External factors like globalization, climate change, loss of biodiversity and exploitation of the bat habitats by urbanization increase contact between humans and bats, and there is an ongoing debate about whether these factors increase the risk of spillover of pathogens to humans [4, 5]. A long-term co-evolution of bats and viruses is assumed, resulting in an adapted immune system and low susceptibility of bats to infection with these pathogens and to the development of symptomatic diseases [6]. This ability of the bat immune system to control the persisting viruses without developing diseases is an interesting research field in terms of immunology [7]. As a result, research on bats in general is an interdisciplinary field that may combine different research questions in One Health, e.g. regarding ecology, zoology and environmental sciences with virology, microbiology, molecular biology, immunology and public health-related issues.

Bats are social animals and live in colonies in different roosting sites including caves. Caves are known to offer largest known mammalian congregations, sometimes exceeding hundreds of thousands of bats roosting in a single location. In these dwellings, bats of different species often roost sympatrically. Although these different species usually occupy different locations inside a cave roost and are essentially separated, they have closer contact during inflight and outflight and also share aerosols. This may facilitate inter-species transmission of viruses and increase the general prevalence and persistence of viruses in these mammals [8]. The interaction between ecological factors and virological investigations is therefore an important part of the interdisciplinary research.

With more than 30 species, bats represent one third of the mammalian species inhabiting the island of Sri Lanka [9]. They contribute to the agro-ecosystems by pollination, seed dispersal and insect control, which renders them particularly interesting for ecological research and conservation biology [9, 10]. Although the general ecology of bats in Sri Lanka is well studied, so far only few scientific studies on their role as viral reservoirs have been published [9, 11– 14]. With this case study, we present the establishment of an interdisciplinary field study on bat species inhabiting Wavul Galge cave (Koslanda, Sri Lanka). In three individual sampling sessions in 2018 and 2019, catching and sampling of bats were optimized in terms of ecological, virological and molecular biological research aspects, personal safety of the research team and reduction of harm and stress for the bats during the sampling process. The planning and conducting of the sampling sessions were based on specific knowledge by experienced zoologists regarding bat behavior, capturing and handling, in combination with the requirements of high-quality sampling for molecular biological and virological analyses. A few guidelines for the catching and handling of bats are available, providing the groundwork for the preparation of sampling sessions [15–17]. However, the field study conditions strongly depend on the environmental circumstances and the respective scientific question that has to be answered. The number of captured bats and the extent of different samples taken per bat may differ. This case study describes the experience we gained throughout this process of optimizing our sampling sessions on bat research. The methods and insights that we describe here may help other research teams when engaging in bat field work also in other countries and when preparing and deciding on the different available sampling strategies.

## Methods

The sampling of bats was carried out in Wavul Galge (Koslanda, Sri Lanka, 6°41’50.2”N 81°03’51.0”E), an underground cave in the interior of Sri Lanka (Figure 1) where bats of the genera *Miniopterus, Hipposideros, Rhinolophus* and *Rousettus* roost sympatrically.

**Figure 1:**
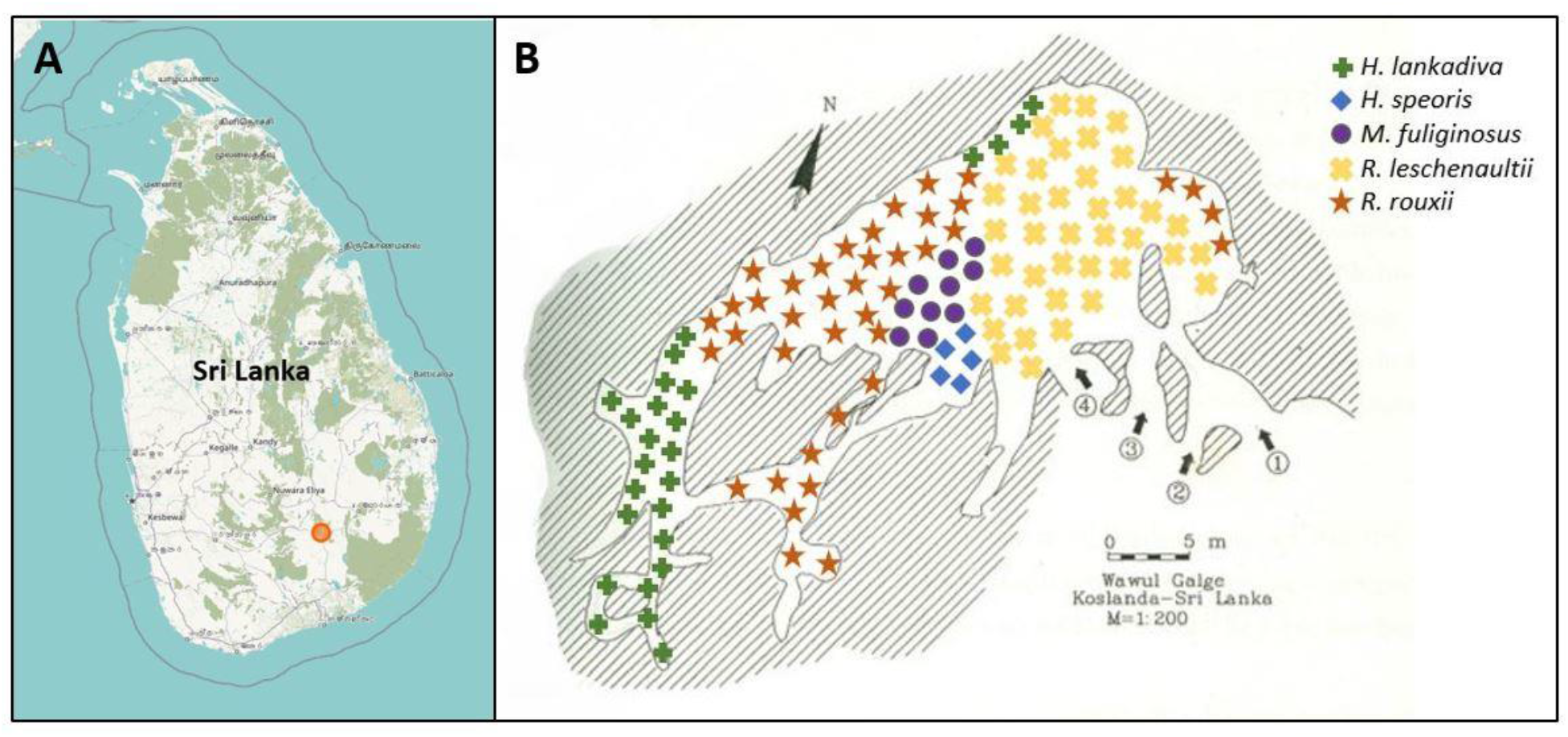
A: Location of Wavul Galge cave (marked in red) on the island of Sri Lanka (© OpenStreetMap in accordance with the Open Data Commons Open Database License). B: Schematic plan of the underground cave Wavul Galge, Sri Lanka. The roosting locations of the different bat species in the cave and the four entrances to the cave (1–4) are shown in the map.

Sampling permission was issued by the Department of Wildlife Conservation, Sri Lanka (permit No. WL/3/2/05/18, issued on 10 January 2018) and the sampling was conducted in accordance with the guidelines and regulations of the Fauna and Flora Protection Ordinance, Sri Lanka.

A total of three sessions for sampling bats were performed in March 2018 (03/18), July 2018 (07/18) and January 2019 (01/19) in order to cover different seasons of the year. During these sampling sessions, we optimized our sampling strategy stepwise and compared different methods and materials. Cost–benefit considerations were made based on stress exposure to the bats, safety precautions for the research team and practicability.

An overview of equipment and material used for conducting the bat samplings is given in the supplementary material.

### Personal protective measures

Adequate personal protective equipment (PPE) was used at all times during the catching and sampling of the bats, both for the protection of the sampling team (to prevent zoonoses) but also to protect the bats (risk of anthropozoonoses).

During the bat catching procedure, head coverings (baseball cap, hats or comparable) and FFP3 masks were worn as protection from bat droppings and aerosols. Solid work gloves were used to transfer the bats from the nets to the bat holding bags.

During the sampling procedure of the bats, different gloves were worn to allow for a more flexible handling of the bats. Instead of rigid work gloves, a layer of conventional lab gloves followed by polyurethane-coated protection gloves were worn. Another layer of lab gloves was used on top to be easily cleaned or changed in the case of contamination.

### Catching procedure

For the catching of bats, either home-made hand nets or a harp trap (Bat Conservation and Management Inc., Carlisle, Pennsylvania, USA) were used and their feasibilities were compared. Furthermore, we compared catching and sampling of bats either in evening sessions at dusk when they left the cave or in morning sessions at dawn when bats returned to the cave. Captured bats were kept in holding bags until further processing; for this a sampling tent was set up near the cave entry. Bats were always released directly in front of the cave after sampling was finished. The catching procedure is visualized in Figure 2.

**Figure 2:**
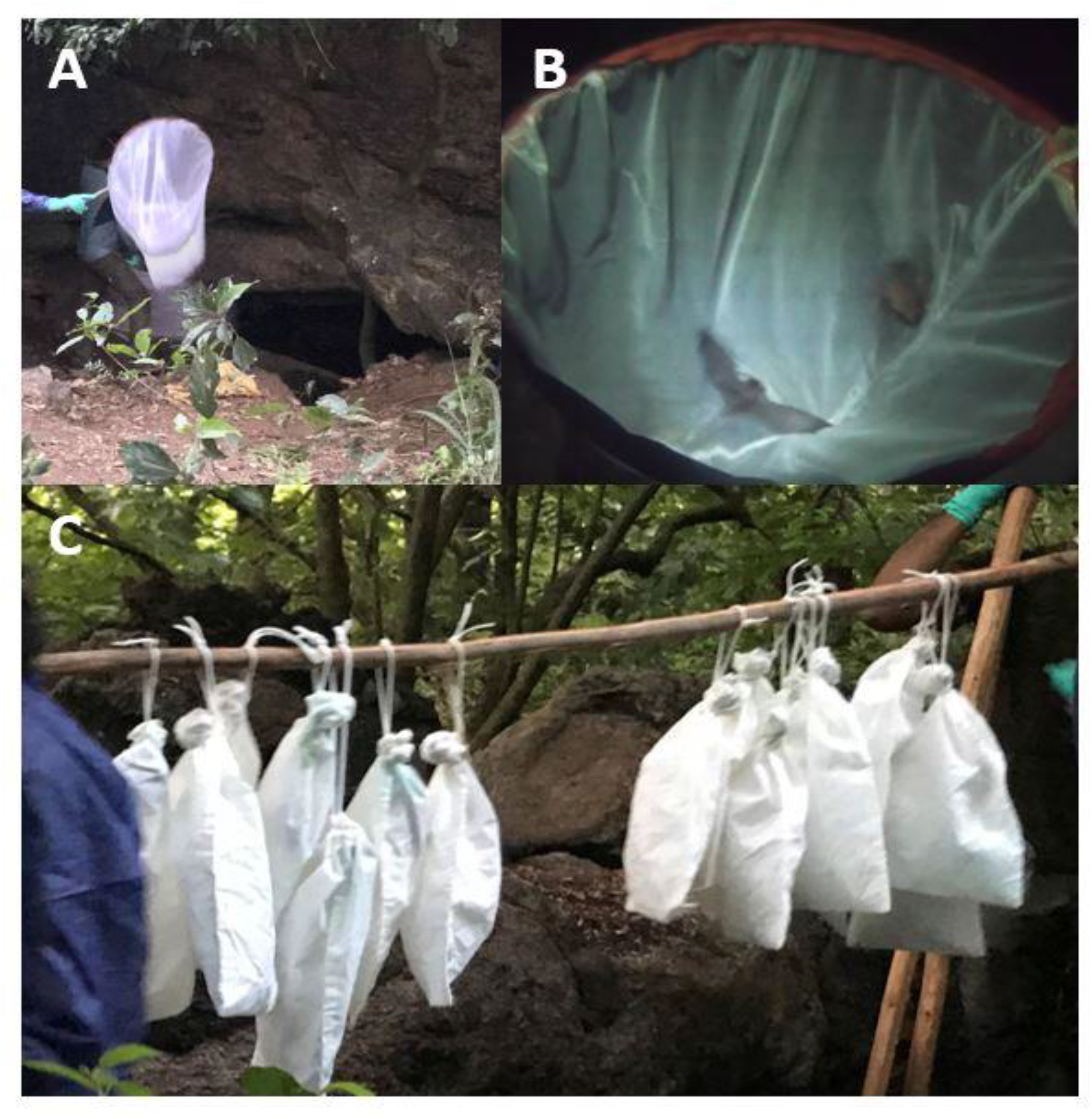
Steps of the bat catching procedure. A: Catching of bats in front of the underground cave Wavul Galge using a hand net. B: Two bats trapped in a hand net. C: Transport of bats in holding bags to the tent for taking samples. © Beate Becker-Ziaja, Therese Muzeniek

### Sampling of bats and documentation

To every individual bat captured an internal number was assigned and basic parameters were assessed and documented as listed in Table 1.

**Table 1:**
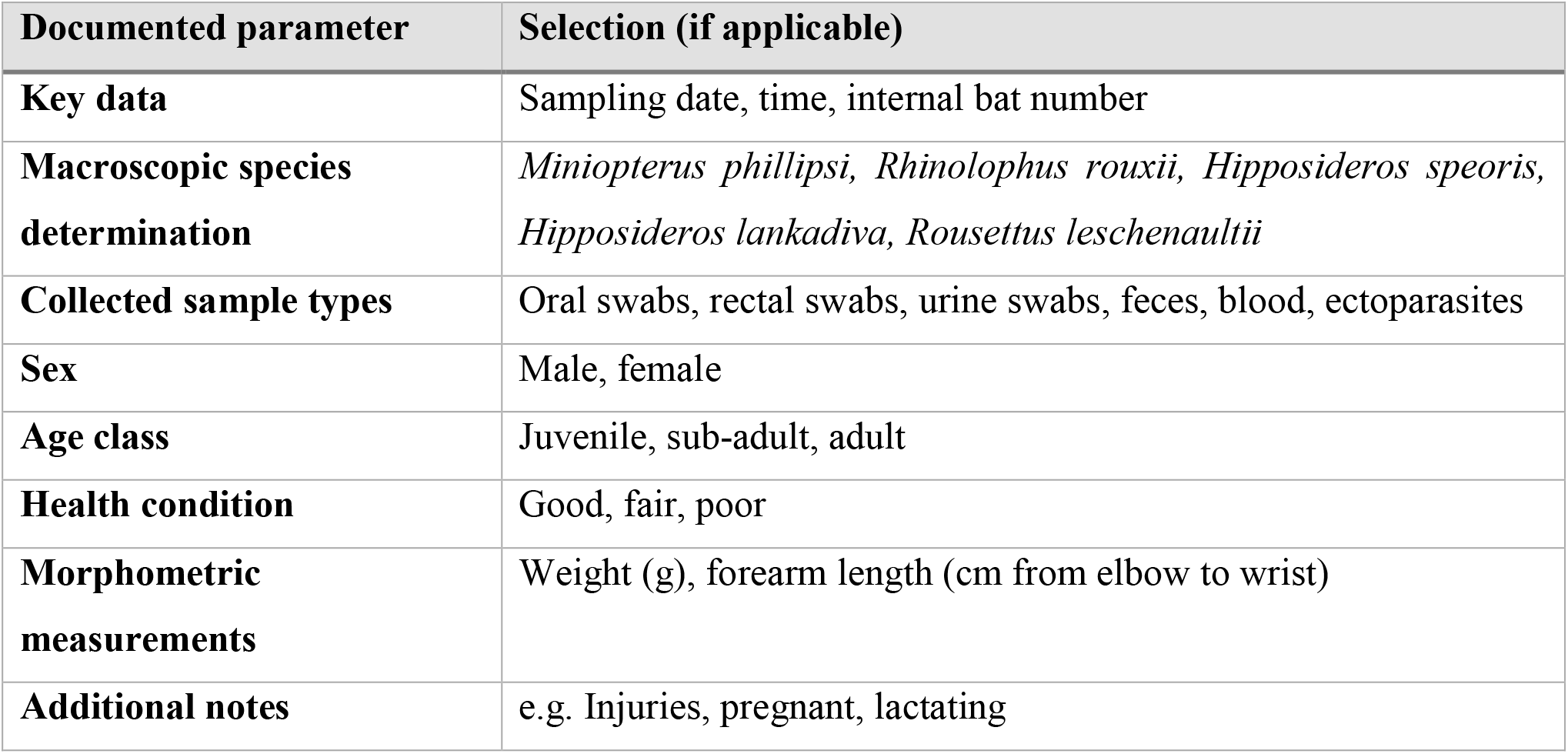
Parameters recorded during the sampling of bats from Wavul Galge cave, Sri Lanka.

According to the issued sampling permission, oral swabs, rectal swabs, urine swabs, fecal pellets, blood samples and ectoparasites were collected from the animals if possible.

For swabbing, we compared sterile Minitip FLOQSwabs® (Copan Diagnostics, Murrieta, CA, USA) with CleanFoam® swabs (ITW Texwipe, Kernersville, NC, USA) which were not sterile but had to be autoclaved in advance. FLOQSwabs® were available in one size and were used during the 03/18 sampling session for taking all kinds of swabs from the bats. CleanFoam® swabs of different sizes and shapes were used during the 07/18 and 01/19 sampling sessions. For taking oral swabs and urine swabs, CleanFoam® round swabs with a diameter of 3.8 mm were used. Rectal swabs were taken by using CleanFoam® spear-shaped swabs with a maximum diameter of 2.5 mm (for more details see supplementary material).

Suitable swabs were selected based on their handiness during sampling and also with regard to the subsequent laboratory processing of the samples. The shapes of the three described swab types are compared in Figure 3.

**Figure 3:**
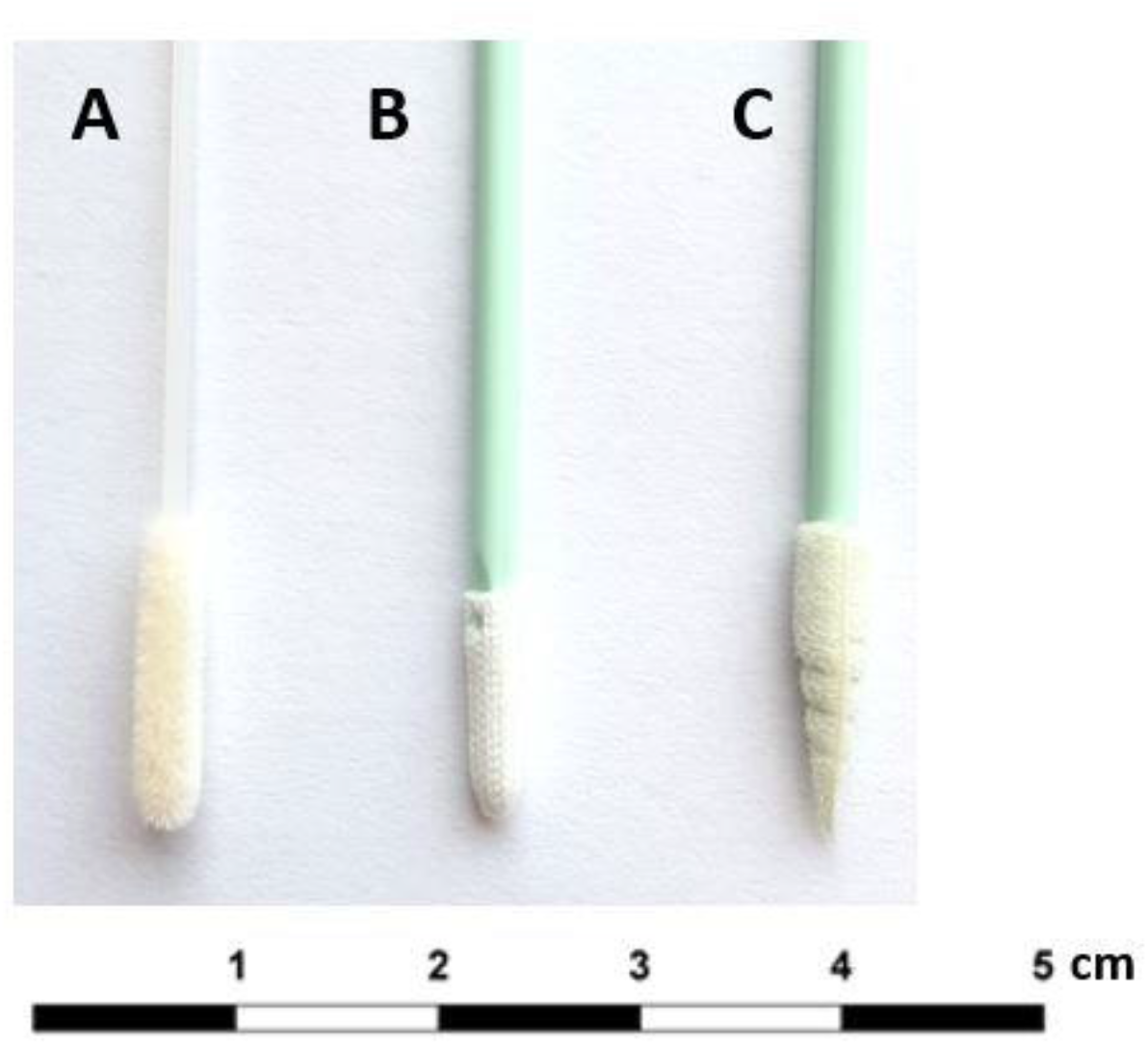
Comparison of different swab types used during the bat sampling sessions. A: FLOQSwabs® used during the 03/18 bat sampling session for all sample types. B: CleanFoam® round swab used during the 07/18 and 01/19 bat sampling sessions for taking oral swabs and urine swabs. C: CleanFoam® spear-shaped swab used during the 07/18 and 01/19 bat sampling sessions for taking rectal swabs.

Fecal pellets were collected with forceps from the bat holding bags if available. Forceps were cleaned afterwards and wiped using ethanol. Ectoparasites were also collected with forceps directly from the bat fur if possible. For taking blood samples, small sterile cannulas (Sterican 30G, 0.3×12mm, B. Braun, Melsungen, Germany) were used. In the first two sampling sessions 03/18 and 07/18, conventional glass capillaries were used for sample collection, stored in 2 mL screw cap microtubes (Sarstedt Inc., Nümbrecht, Germany) and frozen for transportation. In the 01/19 sampling session, blood was collected on Whatman® protein saver cards, stored and transported in plastic bags together with silica gel bags at room temperature.

Examples of the bat sampling procedure are given in Figure 4.

**Figure 4:**
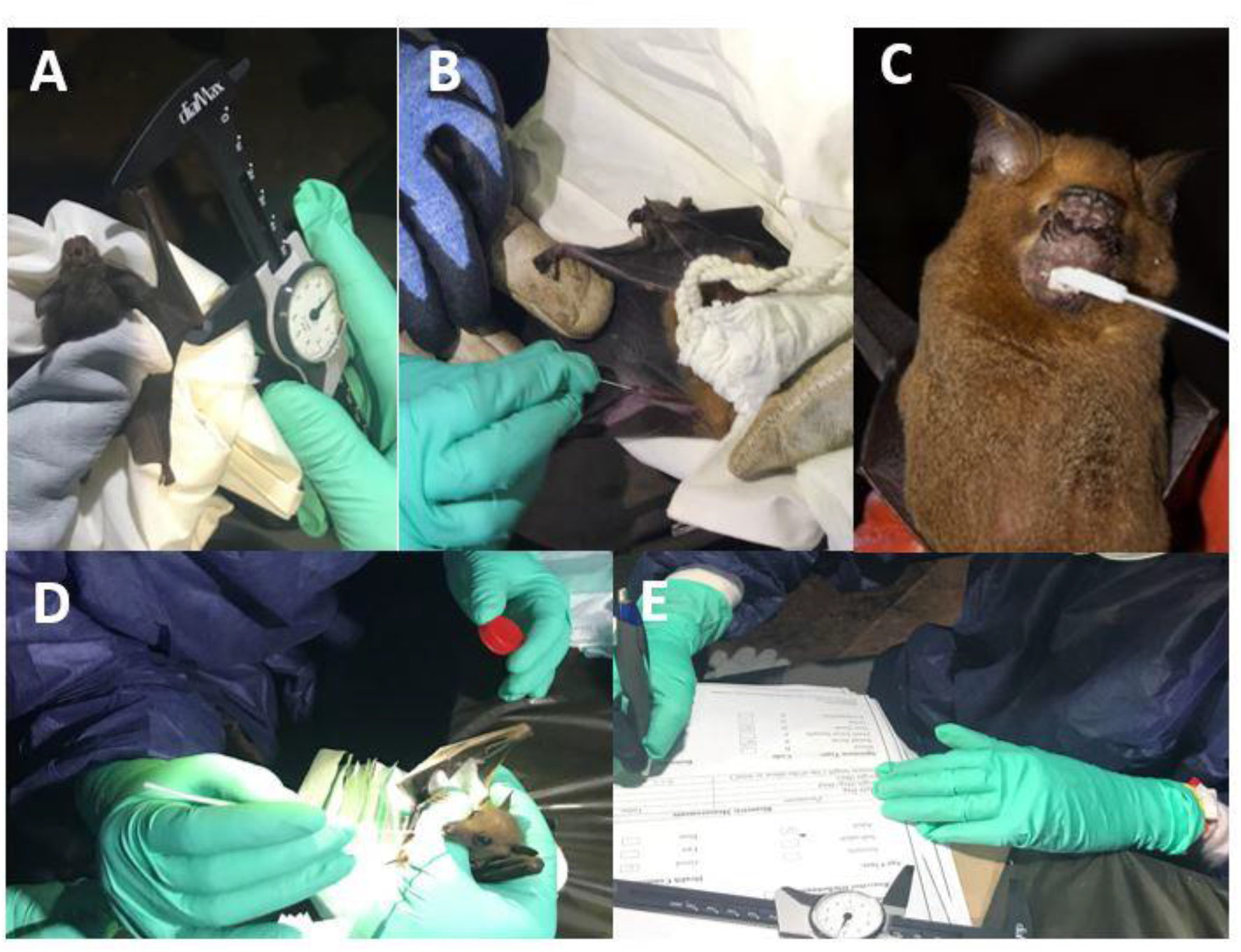
Examples of the bat sampling procedure. A: Taking forearm length measurements of an *R. rouxii* bat. B: Drawing blood from an *H. speoris* bat by venipuncture. C: Oral swabbing of an *H. speoris* bat. D: Oral swabbing of an *R. leschenaultii* bat. E: Documenting the taken measurements on the record sheet. © Beate Becker-Ziaja, Sahan Siriwardana

All collected sample types described above were stored in 2 mL screw cap microtubes and stored without any additives for transportation. During the two first sampling sessions in 03/18 and 07/18, Va-Q-Tcon cooling boxes (va-Q-tec, Würzburg, Germany) with -80°C cooling packs were used for sample storage and transport. In the 01/19 sampling sessions, a dry shipper VOYAGEUR Plus (Cryopal, Bussy Saint Georges, France) with absorbed liquid nitrogen was used for storage and transportation. For this purpose, samples were stored in a nylon stocking and collected samples were separated with a knot to differentiate between the samples per bat. Filled nylon socks were put into the dry shipper for storage. The sample storage is shown in Figure 5.

**Figure 5:**
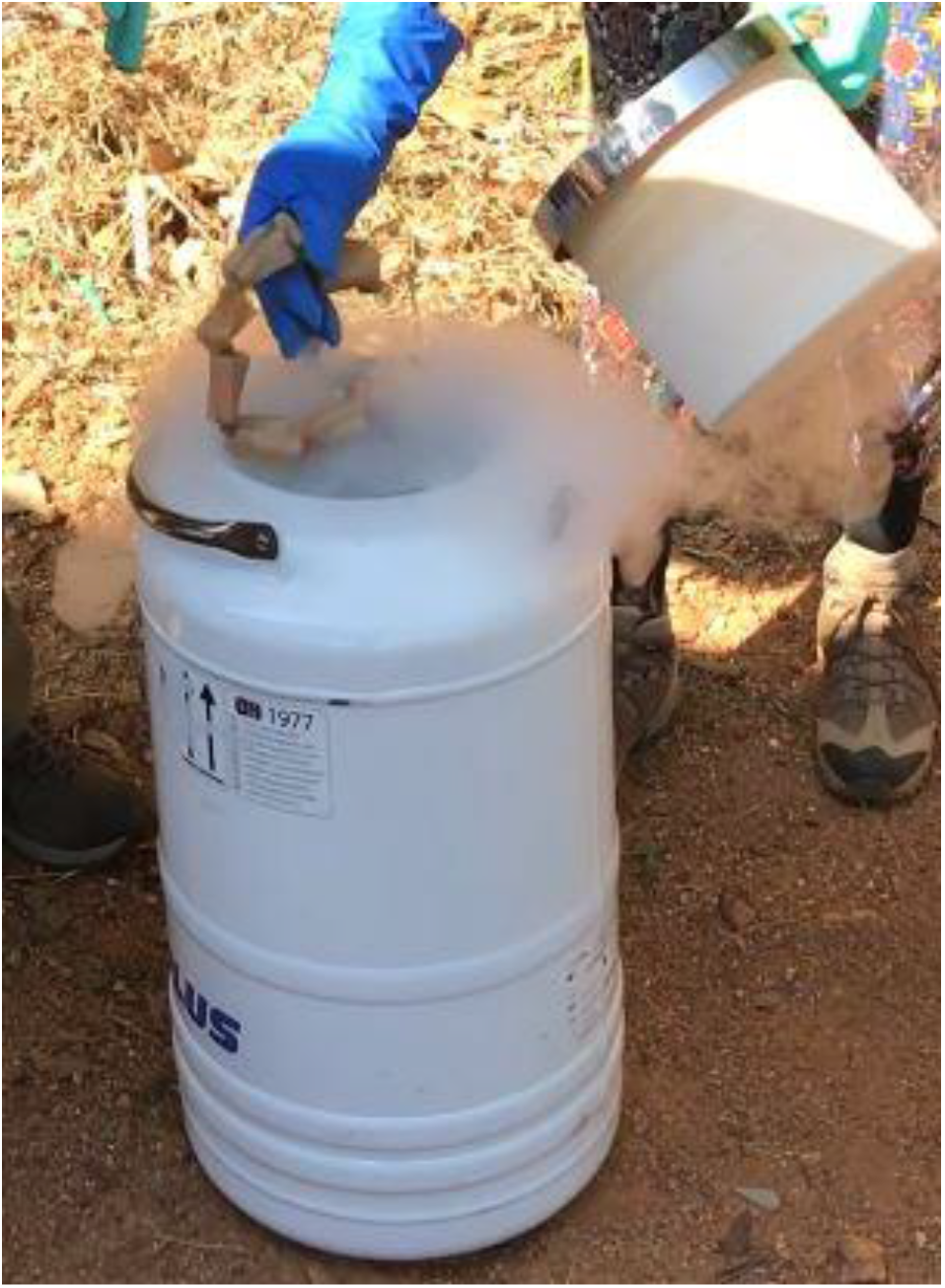
Storing of collected bat samples in a dry shipper by using nylon socks. © Therese Muzeniek

The collected samples were subsequently handled in laboratories of the biosafety level 2 and processed with regards to research questions from zoology, ecology, virology, microbiology, molecular biology, immunology and public health-related issues.

## Results

In a total of three sampling sessions, different bat species from Wavul Galge cave, Sri Lanka, were sampled, while the catching and sampling strategy was optimized stepwise. Table 2 gives an overview of the three sampling sessions including basic conditions and total number of sampled bats. Table 3 gives a more detailed overview of samples collected per bat species.

**Table 2:**
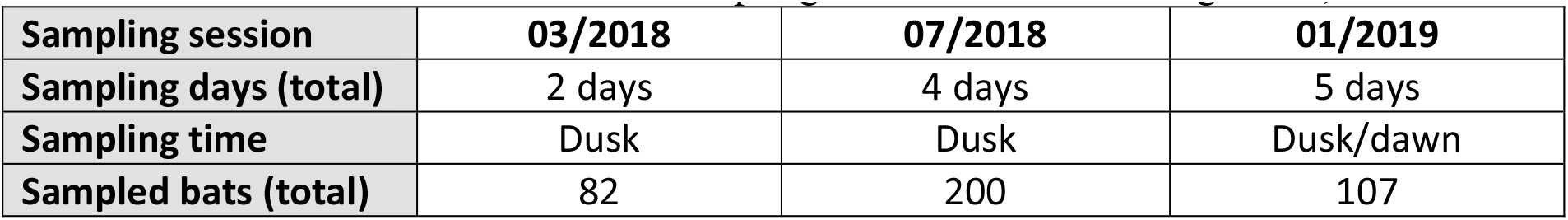
Overview of the conducted bat sampling sessions at Wavul Galge cave, Sri Lanka.

| Sampling session | 03/2018 | 07/2018 | 01/2019 |
| --- | --- | --- | --- |
| Sampling days (total) | 2 days | 4 days | 5 days |
| Sampling time | Dusk | Dusk | Dusk/dawn |
| Sampled bats (total) | 82 | 200 | 107 |

**Table 3:** Overview of the three conducted sampling sessions, including number of collected samples per bat species.

|  |  | Oral swabs | Rectal swabs | Feces | Urine swabs | Blood | Ectoparasites |
| --- | --- | --- | --- | --- | --- | --- | --- |
| ***Hipposideros* spp.** | **Total (bats)** |  |  |  |  |  |  |
| 03/2018 | **3** | 3 | 3 | 0 | 2 | 0 | 0 |
| 07/2018 | **1** | 1 | 0 | 0 | 0 | 0 | 0 |
| 01/2019 | **22** | 22 | 16 | 7 | 6 | 21 | 3 |
| ***M. phillipsi*** | **Total (bats)** |  |  |  |  |  |  |
| 03/2018 | **3** | 3 | 3 | 0 | 0 | 0 | 2 |
| 07/2018 | **188** | 188 | 116 | 77 | 102 | 10 | 10 |
| 01/2019 | **31** | 31 | 4 | 27 | 11 | 23 | 10 |
| ***R. leschenaultii*** | **Total (bats)** |  |  |  |  |  |  |
| 03/2018 | **9** | 9 | 9 | 2 | 2 | 1 | 5 |
| 07/2018 | **11** | 11 | 11 | 0 | 2 | 3 | 5 |
| 01/2019 | **20** | 20 | 16 | 3 | 6 | 18 | 10 |
| ***R. rouxii*** | **Total (bats)** |  |  |  |  |  |  |
| 03/2018 | **67** | 67 | 60 | 8 | 6 | 0 | 10 |
| 07/2018 | **0** | 0 | 0 | 0 | 0 | 0 | 0 |
| 01/2019 | **34** | 34 | 16 | 17 | 6 | 23 | 10 |

As shown in the overview in Table 2, the total days of bat sampling were extended with each sampling session. In doing so, our main goal was not to increase the number of sampled bats; on the contrary, we even reduced the total number of bats during the last visit (Table 3). Instead, we focused on extended sampling including blood samples and collecting feces pellets and ectoparasites, which took considerably more time compared to the other samplings.

In addition, we aimed to balance the number of captured bats per species, which was optimized in the last sampling session in 01/2019. While the use of a harp trap for bat sampling was tested once during the first sampling session, capturing of bats was generally realized using hand nets at the entrances of the cave. This catching procedure also facilitated to control the number of captured individuals per bat species.

### Sampling procedure and documentation

For thorough documentation of the bat samplings, we included different parameters as specified in the methods section (Table 1). These parameters were mainly recorded to cover the ecological and epidemiological aspects of the field study and to obtain an overview on the age, sex and species distribution at different sampling time points. Furthermore, if viruses were identified in subsequent molecular analyses, an epidemiological correlation of the recorded attributes to the positive findings was attempted.

The most suitable sampling material was selected based on different criteria. For swabs the main factors were sterility, absorbing capacity, size and shape. The tested CleanFoam® swabs (round shape) appeared to be most suitable for taking oral swab samples, while the CleanFoam® swabs (mini spear shape) facilitated taking rectal swabs also from smaller bat species like *M. phillipsi*.

For the taking of blood samples, we selected a needle diameter applicable on the smallest bat species which was also suitable for taking blood from larger bats. Although blood collection on filter paper was rather unwieldy compared to using the capillary system, it revealed substantial advantages regarding sample storage, transportation and subsequent sample preparation for lab analyses. Therefore, this method was selected to be the preferred and most suitable one for taking blood.

## Discussion

### Field studies for zoological and molecular biological bat research

With the field studies described, we aimed to capture and sample bats inhabiting Wavul Galge cave in the interior of Sri Lanka. Our goal was to optimize the sampling procedures covering interdisciplinary One Health questions from the fields of zoology, ecology, virology, microbiology, molecular biology, and immunology while minimizing the disturbance to the natural habitat of the bat populations. Sampling was also optimized to increase the safety of the research team. For this purpose, suitable PPE was available. Also, prophylactic measures were considered, including full rabies vaccination and knowledge of hospitals located nearby in case of emergencies. In order to protect the bats from anthropozoonoses, the research team was allowed to work at the cave only in good health condition.

At the same time, we optimized the processes of capturing bats and their sampling in a way to obtain optimal sample quality with minimized stress for the animals and maximum safety of the research team.

### Optimizing the catching of bats

In general, we observed that a smaller number of bats captured per day was beneficial as it reduced the individuals’ total time spent in the holding bags. In return, the number of total days per sampling session was increased. With the low sampling throughput we optimized for our future sampling sessions, the use of hand nets for bat catching appeared to be sufficient.

Advanced sampling equipment would be recommended if a higher number of bats was desired, while a harp trap may be generally advantageous over mist nets regarding the risk of injuring bats that are trapped inside the nets. Mist nets should not be used closer to the entrances of the cave when there is peak exodus of bats, as this may result in entanglement of a large number of bats in the net while releasing them within a short period of time is not feasible.

When planning the duration of the sampling session and the maximum number of bats sampled per day, other parameters like the requested sample types (swabs, blood samples or ectoparasites), the size of the research team and the general sampling conditions (morning, evening and season) have to be considered as they all influence the processing time per bat. Depending on the samples taken, the processing of one bat usually took between five and ten minutes.

During the first two sampling sessions, only evening sessions were performed as the setup of the sampling station and preparation for the catching was facilitated during daytime. The bat catching itself was easy to coordinate, as the different bat species leave the cave in a more or less specific order. Also, sampling itself was more convenient to endure for hours, as temperatures and humidity decrease during the night.

With the last sampling session in 01/2019, we set up sampling in the early morning instead of the evening. Despite the above-mentioned advantages of evening sessions, we observed a number of advantages of morning sessions. The main advantages were observed in the bats as they returned to the cave in the morning, satiated after feeding and calmer compared to the evening. As a result, it was easier to keep the bats in the holding bags until processing and the handling seemed to be less stressful for the bats. When catching bats in the morning, they were more likely to defecate into their holding bags. Thus, the number of feces samples was increased and fewer rectal swab samples had to be taken. Drawing blood especially from small bat species like *Miniopterus phillipsi* was notably easier in the morning sessions using daylight than in evening sessions when only artificial light and head torches were available. The dependence on artificial light was a general disadvantage of the evening sessions, as it attracted numerous insects and at times other wild animals like elephants passing nearby, forcing us to move the bat sampling tent several times to other locations. Therefore, sampling in morning sessions appeared to be safer for the research team, also with regard to personal protection. Due to the rising temperatures over the course of the day, the bat processing was terminated around 10 am to prevent the bats from remaining captured in holding bags too long during the heat of the day. When releasing the bats after processing in the morning, special attention had to be taken to release the bats inside the cave in order to facilitate their orientation and to ensure bats fly directly into the cave. If bats do not enter the cave and circle around the cave in the morning, they could be easy prey for avian predators.

Since it was aimed at carrying out bat research focusing on One Health, comprising ecological aspects and virological questions concurrently, different approaches had to be considered for the field studies. A general overview of the bat species inhabiting the cave including some ecological characteristics that were discovered during these and previous sampling sessions is given in Table 4.

**Table 4:** Overview of some ecological characteristics of the five bat species inhabiting Wavul Galge cave, Sri Lanka, as observed or confirmed during the three bat sampling sessions

|  | *Hipposideros lankadiva* | *Hipposideros speoris* | *Miniopterus phillipsi* | *Rousettus leschenaultii* | *Rhinolophus rouxii* |
| --- | --- | --- | --- | --- | --- |
| **Nutrition** | Insectivorous | Insectivorous | Insectivorous | Frugivorous | Insectivorous |
| **Outflight order** | 2<sup>nd</sup> | 2<sup>nd</sup> | 1<sup>st</sup> | 3<sup>rd</sup> | 1<sup>st</sup> |
| **Reproduction cycles** | March – May | September – October | July – August | April, September | September – October |
| **Migration** | No | No | Yes | No | Yes |

For virus discovery in the bat samples and comparing the virus prevalences among each other, a representative number of individuals per bat species seemed to be useful. However, the proportions of the bat species are not evenly distributed and fluctuate significantly over the year, which is of interest to the ecological research on this sympatric bat colony. For example, during the sampling session in 07/2018 we recognized a peak in *M. phillipsi* bats inhabiting the cave. As was discovered earlier, this bat species uses Wavul Galge as pre-maternity cave during this time of the year (compare Table 4).

Field studies during this time can be of interest under ecological aspects but as well for virological questions, as a number of bats from different surrounding satellite colonies gather and mix in this cave. It is interesting to investigate the virus prevalence and shedding pattern at this time compared to other periods of the year. However, it should be considered that a high number of these bats are pregnant at that time point and therefore should not be stressed, for instance by taking blood.

These examples show that zoological aspects and molecular biological questions have to be balanced thoroughly when preparing the respective field studies. The expertise and experience of zoologists and ecologists are important contributions to virological research in the field. In the following, we discuss the molecular biological requirements that influenced the bat sampling process.

### Bat sampling for subsequent molecular biological and virological analyses

With the samples taken from bats, it was intended to perform different molecular biological and virological investigations including PCR, metagenomic NGS, virus isolation in cell culture and serological analyses. For this spectrum of analyses, we aimed to take different sample types in optimal quality. For successful virus isolation, intact virus particles are required. Furthermore, intact nucleic acids with low degradation are needed for PCR and NGS analyses. Both the virus particles and nucleic acids are susceptible to high temperatures and may degrade quickly. A continuous cold chain was therefore essential for the sample quality and successful laboratory analysis, especially in a tropical country like Sri Lanka. We observed that even professional cooling bags could not maintain an adequate temperature for a longer period of time. A suitable solution was therefore the use of a dry shipper that maintained the temperature longer. This shipper contained material adsorbing liquid nitrogen, and the cooling was achieved by its gradual evaporation into the storage area of the shipper. By this, an adequate cooling temperature could be maintained for several weeks. In addition, in accordance with current IATA regulations, appropriate dry shippers are suitable for transport of the samples via airplane, facilitating the sample logistics.

Due to the limited size of the available dry shipper, its limited capacity for sample storage and transportation had to be considered when planning the respective field trips. Therefore, a beneficial solution for taking blood samples was to collect drops on protein saver cards. These were stored in a single plastic bag per card and transported at room temperature. It was important to avoid humidification of the dried blood; for this we used desiccants like silica gel sachets in each bag.

Other samples such as oral swabs, rectal swabs, urine and feces were snap-frozen within the dry shipper without any additives. Liquid was added later during the preparation process of the samples in the laboratory. In terms of molecular biological analyses, this method did not impair the outcome of the analyses as we were able to detect different viruses via PCR and in NGS analyses [11, 12, 18]. The presence of intact virus particles has not been tested yet, and virus isolation in cell culture will be performed in future analyses. However, samples may also be stored in suitable viral transport medium or PBS in order to maintain virus integrity. Care must be taken to use a medium suitable for the variety of subsequent analyses.

It is a common technique to take wing punches of the sampled bats for species identification purposes [19]. However, it was discovered that oral swabs were also suitable for the analysis based on cytochrome b PCR and sequencing [20]. By this, it was possible to reduce invasive bat sampling to a minimum.

Concerning different kinds of tested swabs, we observed that a less absorbing material was beneficial especially for the subsequent sample preparation. For the tested FLOQSwabs®, additional 300 µL of liquid had to be added in order to retain sufficient volume for Nucleic Acid extraction. In contrast, the preferred CleanFoam® swabs absorbed the liquid samples (urine, saliva etc.) but clearly less of the liquid in the subsequent processing. In addition, the stable handle and special forms of the different CleanFoam® swabs facilitated sample taking, especially for small bat species. However, other swab brands may be suitable depending on the respective sampling setup. It is important to consider the sample material of the swabs, as any organic material (e.g. cotton) would interfere with NGS analyses by increasing the sequencing background [19]. Furthermore, swabs should be sterile or autoclavable in order to allow for samples free of contamination. But also vice versa, the material was selected in order to be harmless for the health of the sampled bats.

## Conclusion

We performed three individual bat sampling sessions to optimize sampling of multiple bat species roosting sympatrically in Wavul Galge cave, Sri Lanka. The samplings took place at different times of the year in order to adjust to different circumstances regarding bat population dynamics and colony size, breeding seasons and environment variables such as temperature and humidity. We compared different sampling setups and optimized the total duration of a sampling session, the sampling time point, the sampling itself and sample logistics in terms of storage and cold chain. All aspects were optimized in order to cause as little stress as possible for the sampled bats, to increase the safety of the research team and to obtain optimal sample output at the same time. The experiences gained during this research will help to perform frequent sampling sessions in the future and to monitor the wellbeing and conservation of this colony. The long-term observation will help to better understand zoological questions such as population dynamics and interaction of the different species. Molecular biological analyses and detection of viral shedding will help to link these ecological aspects to virological and public health-relevant questions. We expect that our observations and the subsequent optimization process for the sampling sessions will help other teams to plan or improve comparable bat sampling experiments in the context of One Health.

## Acknowledgement

The authors are grateful to Ursula Erikli for copy-editing and to Marica Grossegesse and to Nicole Kromarek for supporting the organization and realization of the first bat sampling session at Wavul Galge cave, Sri Lanka. Further we thank the Department of Wildlife Conservation, Sri Lanka for granting the research permit to carry out the field work and for granting the export permission of samples.

