## Supplemental Material for "One Health Approach for the sampling of different bat species living in a sympatric colony"

### Supplementary Table

Table 1: Overview of material that was used as PPE and for catching and sampling of bats during the three conducted bat sampling sessions in Wavul Galge, Sri Lanka. If applicable, manufacturer and article details of the used equipment is indicated, but may be replaced by comparable articles.

| **Personal protective equipment (PPE)** | | |
|---|---|---|
| **Description** | **Application** | **Manufacturer, article details (if applicable)** |
| Baseball cap or hat | Head protection from bat droppings | Any suitable head protection |
| FFP3 mask | Protection from aerosols | 3M Company, Saint Paul, Minnesota, USA Aura™ 9330+ (Article 3M-ID 7000088812) |
| Work gloves | Transfer of bats after catching | Any, available in construction markets |
| Lab gloves | Handling / Sampling of bats | MICROFLEX® NeoTouch™ (Article 25-201) |
| PU cut protection gloves | Handling / Sampling of bats | Engelbert Strauss, GmbH & Co. KG, Biebergemünd, Germany (Article #7616407) |
| **Bat Catching** | | |
| **Description** | **Application** | **Manufacturer, article details (if applicable)** |
| G7 Harp Trap | Trapping of bats | Bat Conservation and Management Inc., Carlisle, Pennsylvania, USA |
| Hand net | Trapping of bats | Self-made, suitable nets are also available in shops for angling accessories |
| Bat holding bats | Trapping of bats | NHBS GmbH, Bonn, Germany (Article #234513) |
| **Bat sampling** | | |
| **Description** | **Application** | **Manufacturer, article details (if applicable)** |
| Minitip FLOQSwabs® | Suitable for oral swabs | Copan Diagnostics, CA, USA (Article 501CS01) |
| CleanFoam® Swabs, round shape, 3.8 mm | Preferred swab for taking oral and urine swabs | ITW Texwipe, Kernersville, NC, USA (Article TX741B) |
| CleanFoam® Swabs, spear shape, 2.5 mm max | Preferred swab for taking rectal swabs | ITW Texwipe, Kernersville, NC, USA (Article TX751B) |
| dialMax Vernier Dial Caliper | Taking forearm length measurements of bats | Wiha Werkzeuge GmbH, Schonach, Germany (Article 27082) |
| Light-Line Spring Scale | Weighing of bats | PESOLA Präzisionswaagen AG, Schindellegi, Switzerland (Article 10050) |
| Forceps | Collection of fecal pellets, collection of ectoparasites | Bürkle GmbH, Bad Bellingen, Germany Article (5386-0300) |
| Sterile cannulas (Sterican 30G, 0.3x12mm) | Taking blood from bats | B. Braun, Melsungen, Germany (Article 4656300) |
| Whatman® protein saver cards | Storage of collected blood | Cytiva, Marlborough, Massachusetts, USA (Article 10531018) |
| 2 ml screw cap mini tubes | For storage of collected samples, sterile | Sarstedt AG & Co. KG, Nümbrecht, Germany (Article 72.694.306) |
| Va-Q-Tcon Cooling Boxes | Short-term storage of samples (1 – 2 days) | va-Q-tec AG, Würzburg, Germany (Article BC000102) |
| Dry shipper VOYAGEUR Plus | Freezing and long-term storage of samples | Cryopal, Bussy Saint Georges, France (Article VOYAGEUR20-2) |
| Nylon socks | Facilitated storage of samples in the dry shipper | Any available socks |
